# *Brassica napus* genes *Rlm4* and *Rlm7*, conferring resistance to *Leptosphaeria maculans*, are alleles of the *Rlm9* wall-associated kinase-like resistance locus

**DOI:** 10.1101/2021.12.11.471845

**Authors:** Parham Haddadi, Nicholas J. Larkan, Angela Van de Wouw, Yueqi Zhang, Ting Xiang Neik, Elena Beynon, Philipp Bayer, Dave Edwards, Jacqueline Batley, M. Hossein Borhan

## Abstract

*Brassica napus* (canola/rapeseed) race specific resistance genes against blackleg disease, caused by the ascomycete fungus *Leptosphaeria maculans*, have been commonly used in canola breeding. To date; *LepR3*, *Rlm2* and *Rlm9 R* genes against *L. maculans* have been cloned from *B. napus*. LepR3 and Rlm2 are Receptor Like Proteins (RLP) and the recently reported Rlm9 is a Wall Associated Kinase-Like (WAKL) protein. *Rlm9* located on chromosome A07 is closely linked with *Rlm3, Rlm4, RLm7* genes. Recognition of AvrLm5-9 and AvrLm3 by their corresponding Rlm9 and Rlm3 proteins is masked in the presence of AvrLm4-7. Here we report cloning of *Rlm4* and *Rlm7* by generating genome sequence of the doubled haploid (DH) *B. napus* cv Topas DH16516 introgression lines Topas-*Rlm4* and Topas-*Rlm7*. Candidate *Rlm4* and *Rlm7* genes were identified form the genome sequence and gene structures were determined by mapping RNA-sequence reads, generated from infected cotyledon tissues, to the genome of Topas-*Rlm4* and Topas-*Rlm7*. *Rlm4* and *Rlm7* genomic constructs with their native promoters were transferred into the blackleg susceptible *B. napus* cv Westar N-o-1. Complementation of resistance response in the transgenic Westar:*Rlm4* and Westar:*Rlm7* that were inoculated with *L. maculans* transgenic isolates 2367:*AvrRlm4-7* or 2367:*AvrLm7* confirmed the function of *Rlm4* and *Rlm7* genes. Wild type *L. maculans* isolate 2367 that does not contain *AvrLm4-7* or *AvrLm7*, and transgenic 2367:*AvrLm3* and 2367:*AvrLm5-9* did not induce resistance proving the specificity of *Rlm4* and *Rlm7* response. *Rlm4* and *Rlm7* alleles are also allelic to *Rlm9*. *Rlm4* and *Rlm7* genes encode WAKL proteins. Comparison of highly homologous sequences of *Rlm4* and *Rlm7* with each other and with the sequence of additional alleles, using whole genome sequencing of additional 128 lines, identified a limited number of point mutation located within the predicted extracellular receptor domains.

## Main Text

Plant cell surface receptors are at the forefront of defense against pathogens, involved in pathogen sensing through the detection of conserved molecules named pathogen-associated molecular patterns (PAMP) and highly variable pathogen virulence (effector) proteins (Albert et al., 2019). The recently reported *Brassica napus* (canola, oilseed rape) disease resistance gene *Rlm9* encodes a wall-associated kinase-like (WAKL) receptor which confers race-specific resistance against races of the blackleg pathogen *Leptosphaeria maculans* carrying the corresponding effector gene, *AvrLm5-9* (Larkan et al., 2020). WAKLs are transmembrane proteins which bridge the intracellular kinase domain to the extracellular galacturonan-binding (GUB), WAK and epidermal growth factor (EGF)-like calcium binding domains (Verica and He, 2002).

*Rlm4* and *Rlm7* are located on *B. napus* chromosome A07 and genetically tightly-linked to *Rlm9* (Larkan et al., 2016). The *L. maculans* effectors AvrLm4-7 and AvrLm7 are small, secreted cysteine-rich proteins encoded by a single locus, *AvrLm4-7*. A single amino acid change in AvrLm4-7 masks recognition by Rlm4 without affecting Rlm7 function (Parlange et al., 2009). Here we report the cloning of *Rlm4* and *Rlm7*, both alleles of the *Rlm9* WAKL locus.

Genomic sequencing data was generated for the *B. napus* introgression line Topas-*Rlm4* and the newly-produced Topas-*Rlm7* (produced as described by Larkan et al., 2016 using the *B. napus* variety Roxet as the *Rlm7* donor line) using Illumina HiSeq 2500 (2 x 250bp, 4 lanes). Close to 537 million reads (113.64x coverage) were assembled using SOAPdenovo assembler. Contigs generated from each line were mapped to the *B. napus* reference genome *‘Darmor-bzh’* using Bowtie2 and visualized using CLC. Based on ~1.3 billion RNA sequence reads mapped to the *Rlm3-4-7-9* gene cluster (Figure 1a), *Rlm4* and *Rlm7* genes were determined to be allelic, with each being 8507 bp in length consisting of three exons (1013bp, 126 bp, 1240 bp) and two introns (106 bp and 6022 bp) (Figure 1b). To prove the function of the predicted genes, the entire gene and 5’ intergenic region (1750 bp) for each allele was synthesized (GenScript, USA) and cloned into the plant transformation vector pMDC123, modified to contain the *nosT* terminator sequence downstream of the cloning site (pMDC123-*nosT*), using Gateway cloning technology. *Rlm4* and *Rlm7* genomic constructs were transferred into the blackleg-susceptible *B. napus* doubled-haploid line Westar N-o-1 as previously described (Larkan et al., 2020). Regenerated transgenic (T_0_) plants that survived herbicide selection were screened via droplet digital PCR (ddPCR) to identify lines carrying insertions, then selfed to produce the T_1_ generation. The resulting transgenic lines (13 for *Rlm4*, 17 for *Rlm7*) were initially tested for resistance response using the *L. maculans* isolate v23.1.3 (*avrLm3, AvrLm4-7, avrLm9*) using a standard cotyledon assay (Larkan et al., 2013) with all lines displaying hypersensitive response at the point of infection except for one *Rlm4* line with poor germination (data not shown). Additional ddPCR was conducted to identify homozygous, single insertion events in T_1_ plants, and one plant for each construct was selected and selfed to produce homozygous T_2_ lines for further characterisation (hereafter referred to as Westar:*Rlm4* and Westar:*Rlm7*). Further confirmation was obtained by utilising transgenic *L. maculans* to demonstrate effector-specific activation of resistance conferred by the *Rlm4* and *Rlm7* candidate genes. The *L. maculans* isolate 2367 (*avrLm3, avrLm4-7, avrLm9*) and the transgenic isolates 2367:*AvrLm4-7*, 2367:*AvrLm7* (Larkan et al., 2016) and 2367:*AvrLm5-9* (Ghanbarnia et al., 2018) were used to inoculate the *B. napus* lines Westar N-o-1 (susceptible), Westar:*Rlm4*, Westar:*Rlm7* (this study) and Westar:*Rlm9* (Larkan et al., 2020). Four seedlings of each line were inoculated with each isolate (one inoculation per cotyledon lobe; four per seedling), with the whole experiment being performed in duplicate. No resistance reaction was induced in either Westar:*Rlm4* or Westar:*Rlm7* in response to *AvrLm5-9*. A hypersensitive response was induced in Westar:*Rlm4* only in response to *AvrLm4-7*, while Westar:*Rlm7* responded to both *AvrLm4-7* and *AvrLm7*, as expected, confirming the cloned genomic constructs as *Rlm4* and *Rlm7*, respectively (Figure 1c).

**Figure 1.**
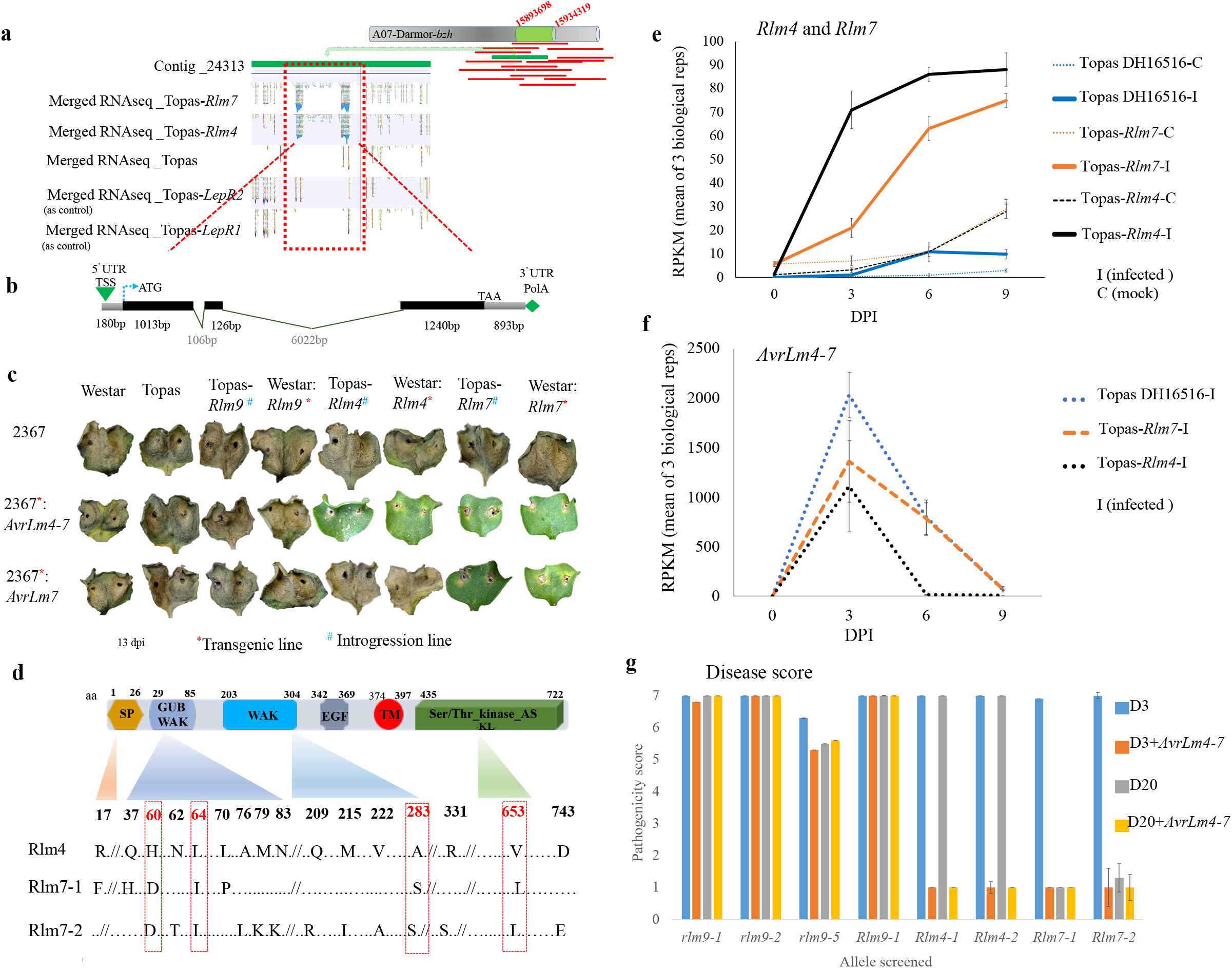
Cloning of two allelic *B. napus*, wall-associated kinase-like (WAKL) genes, *Rlm4* and *Rlm7*. **(a) Identification of *Rlm4* and *Rlm7* through contig walking.** Merged RNA seq from the early stage of infection (3-dpi) revealed the expression of *Rlm4* and *Rlm7* in single gene introgression lines Topas-*Rlm4* and Topas-*Rlm7*. **(b) *Rlm4* and *Rlm7* gene structures**. Determined by merged RNA-sequence reads, were similar and are depicted by exon (as black bar), intron (black line valley) and UTR (gray) using the merged RNA-Seq reads. **(C) Transgenic *B. napus* expressing *Rlm4* and *Rlm7*.** Cotyledons of Westar and Topas (susceptible to blackleg disease), *Topas-Rlm9* and Westar:*Rlm9* (negative controls), Topas-*Rlm4* and Topas-*Rlm7* (positive controls), Westar:*Rlm4* and Westar:*Rlm7* transgenic lines 13 dpi with isolate 2367 (virulent towards *Rlm4, Rlm7 and Rlm9*) and the transgenic isolates 2367:*AvrLm4-7* (avirulent toward *Rlm4* and *Rlm7*) and 2367:*AvrLm7* (avirulent toward *Rlm7*). **(d) Protein domains of Rlm4 and Rlm7.** Protein structure is represented by the signal peptide (SP-yellow bar), extracellular GUB_WAK pectin-binding (dark blue), C-terminal WAK (blue bar) and EGF-like Ca^2+^ (EGF, gray bar), transmembrane (TM, red bar) and an intracellular serine/threonine Protein Kinase (green bar) domains. (**e, f) Expression profile of *Rlm4, Rlm7* and *AvrLm4-7*.** RNA-Seq time course (0, 3, 6 and 9 dpi) analysis showed a significant upregulation of *Rlm4* and *Rlm7* during *L. maculans* infection when compared to the mock. No significant induction was observed in susceptible (*rlm4-7*) line Topas DH16516. (**g) Confirmation of *Rlm4* and *Rlm7* alleles through pathogenicity phenotyping.** *B. napus* lines and cultivars harbouring different susceptible alleles (*rlm9-1, rlm9-2, rlm9-5*) or *Rlm9 (Rlm9-1), Rlm4 (Rlm4-1, Rlm4-2*) or *Rlm7 (Rlm7-1, Rlm7-2*) resistant alleles were screened for resistant (pathogenicity scores <3.0) and susceptible reactions (pathogenicity scores >5.0) using progenitor differential isolates (D3 and D20) as well as genetically modified isolates harbouring *AvrLm4-7* (D3+*AvrLm4-7*, D20+*AvrLm4-7*).

*Rlm4* and *Rlm7* open reading frames are 2,379 bp encoding highly homologues proteins of 792 amino acids (aa). Sequence polymorphism between the two genes is limited to a total of 13 single nucleotide substitutions resulting in four synonymous and nine non-synonymous changes. Interpro predicts Rlm4 and Rlm7 as transmembrane proteins consisting of extracellular WAK_GUB (aa 29-85), WAK (aa 203-314), EGF like calcium binding domains (aa 342-369) and a cytoplasmic kinase domain (aa 435-722) (Figure 1c). The first 26 aa in both proteins are the predicted secretory signal peptide (SP). There are a total of 7 amino acid differences between Rlm4 and Rlm7. Amino acid changes (Rlm4 > Rlm7) are at positions 17 (SP; F:R), 37 (GUB; Q:H), 60 (GUB; H:D), 64 (GUB; L:I), 70 (GUB; L:P), 283 (WAK; A:S) and 653 (Kinase: L:V) (Figure 1d). A further 128 lines/cultivars that had been phenotyped for the presence of *Rlm4* and *Rlm7* using differential isolates were investigated using whole genome sequencing leading to the identification of two additional resistant alleles (*Rlm4-2* and *Rlm7-2*) The ORF sequences of the *Rlm4-1* and *Rlm4-2* alleles were identical, with any polymorphisms limited to the intronic regions of the gene. The *Rlm7-2* allele, identified in the *B. napus* variety Caiman, contained 13 non-synonymous SNPs, however these aa changes do not affect the recognition of Rlm7. The Rlm7-1 and Rlm7-2 proteins differ from Rlm4 in conserved amino acids in the putative extracellular ligand-binding domains, at positions 60, 64 (WAK_GUB) and 283 (WAK), which may explain the variation in recognition specificity between Rlm4 and Rlm7 towards AvrLm7. An additional conserved polymorphism is found in the kinase domain (at position 653), though this is unlikely to be involved in recognition of AvrLm4-7 (Figure 1d).

The Topas-*Rlm4* and Topas-*Rlm7* introgression lines were utilised to monitor the expression of *Rlm4*, *Rlm7* and the corresponding Avr gene *AvrLm4-7* during cotyledon infection by the reference isolate v23.1.3 (RNA processing was as previously described by Haddadi et al., 2019). Expression of *Rlm4* and *Rlm7* increased substantially in both the resistant Topas-*Rlm4* and Topas-*Rlm7* lines in response to infection, though upregulation of *Rlm7* was somewhat less rapid than that of *Rlm4* (23- vs 3-fold increase early in the infection, 3 days post-inoculation (dpi), FDR < 0.001) (Figure 1e). *AvrLm4-7* expression peaked at 3 dpi but rapidly declined at later time points (Figure 1f).

Additional pathology tests using representative *B. napus* lines harbouring both *Rlm4* and *Rlm7* alleles, as well as the *Rlm9* and four of the susceptible alleles, was performed with *L. maculans* isolates transformed with a functional copy of *AvrLm4-7*. Progenitor isolate D3 is virulent towards both *Rlm4* and *Rlm7* whilst progenitor isolate D20 is only virulent towards *Rlm4*. Isolate D3+*AvrLm4-7* resulted in an avirulent reaction on lines harbouring each of the *Rlm4* and *Rlm7* alleles but remained virulent on all other lines as expected. Isolate D20+*AvrLm4-7* resulted in an avirulent reaction on lines harbouring both *Rlm4* alleles but remained virulent on the lines harbouring the susceptible alleles or *Rlm9* and remained avirulent on the *Rlm7* lines as expected (Figure 1g).

The cloning of *Rlm4* and *Rlm7* will provide valuable information for canola breeding programs worldwide. It also increases the number of cloned WAKL-type race-specific *R* genes to four (Saintenac et al. 2018; Larkan et al. 2020; this study), expanding the toolbox for further study of this newly-emerging class of plant *R* genes.

## Author Contributions

NJL and HB conceived and designed the study, PH and HB extracted the *Rlm4-1* and *Rlm7-1* ORFs and designed the transgenic constructs, PH produced and analysed RNA-Seq data, NJL and EB analysed and selected transformants, EB performed all ddPCR, AVdW characterised additional Rlm4 and Rlm7 lines, YZ and TXN performed resequencing and characterisation of additional *Rlm4* and *Rlm7* alleles, PB and DE provided bioinformatic analysis. PH, NJL, AVdW, JB and HB prepared the manuscript.

## Data Availability

All sequence data is submitted to GenBank under the following accession numbers:

BankIt2528103 Rlm4 OL806566
BankIt2528103 Rlm7 OL806567
BankIt2528103 Rlm4_1 OL806568
BankIt2528103 Rlm4_2 OL806569
BankIt2528103 Rlm7_1 OL806570
BankIt2528103 Rlm7_2 OL806571

